# Autotransporter secretion exploits the bacterial actin-homologue

**DOI:** 10.1101/305441

**Authors:** Mahmoud M. Ashawesh, Robert Markus, Christopher N. Penfold, Kim R. Hardie

## Abstract

Bacterial infection of humans, animals and plants relies heavily on secreted proteases that degrade host defences or activate bacterial toxins. The largest family of proteins secreted by Gram-negative pathogenic bacteria, the Autotransporters (ATs), includes key proteolytic virulence factors. There remains uncertainty about the mechanistic steps of the pathway ATs share to exit bacteria, and how it is energetically driven. This study set out to shed light on the AT secretion pathway with the ultimate aim of uncovering novel antimicrobial targets that would be unlikely to trigger the development of resistance mechanisms in bacteria. To do this, two AT virulence factors with distinct proteolytic functions, EspC (secreted from Enteropathogenic *Escherichia coli*) and AaaA (tethered to the extracellular surface of *Pseudomonas aeruginosa*) were chosen. EspC and AaaA were fluorescently labelled using two separate methods to establish the localization patterns of ATs as they are secreted from a bacterial cell. Super resolution microscopy revealed that localization of ATs occurs via a helical route along the bacterial cytoskeleton. In addition to requiring the conserved C-terminal β-barrel translocator domain of the AT, we present the first evidence that secretion is dependent on a dynamic interaction with a structure reliant upon the actin homologue MreB and the Sec translocon. These findings provide a step forward in the mechanistic understanding of the secretion of this widely distributed family of proteins that have pivotal roles in bacterial pathogenesis and conserved structural properties that could serve as novel broad-range antimicrobial targets.

**Significance:** Secreted bacterial proteases facilitate the infection of human, animal and plant hosts by degrading host defences or activating bacterial toxins. The autotransporter family is the largest family of proteins secreted from Gram-negative bacteria, and includes proteolytic virulence factors crucial to bacterial infection. Precisely how autotransporters migrate from the inside to the outside of the cell, and how this movement is energetically driven is a mystery. We demonstrate a spiral pathway of autotransporter secretion, presenting evidence that it involves a dynamic interaction with the actin homologue MreB that comprises the bacterial cytoskeleton. Our findings open the way to unravelling the mechanism of autotransporter secretion and offer the possibility to identify novel antimicrobial targets unlikely to trigger the development of antimicrobial resistance.

## Introduction

Infections caused by Gram-negative bacteria are a major threat to the health of humans, plants and animals. The widely distributed autotransporter (AT) family includes numerous virulence factors that are vital to the pathogenicity of Gram-negative bacteria. ATs bear conserved structural features offering an ideal target for novel broad spectrum antimicrobials. Novel antimicrobial agents are desperately needed because those currently available have become less effective due to the emergence of antibiotic resistance in hospitals and communities (1, 2). Virulence factors are only required during host infection offering targets less likely to trigger the development of resistance mechanisms in the bacteria (3). Since ATs are extracellular they are easily accessible to antimicrobials. Moreover, the AT secretion pathway offers an ideal target for novel antimicrobials (4–7) as it includes targets unique to bacteria that can be either narrow or broad spectrum (8).

Although the main steps of AT secretion have been identified, many mechanistic details remain unresolved and are hotly debated (7, 9, 10). What at first appeared a self-sufficient secretion pathway has since been shown to comprise of multiple components including the inner membrane (IM) Sec translocon, periplasmic chaperones and the outer membrane (OM) β-barrel assembly machinery (BAM) complex or two-membrane spanning complex TAM (Translocation and assembly module). How and where these components interact together to drive secretion remains to be elucidated, and could have a bearing on the overall building of the bacterial cell envelope because of the pleiotropic roles of the machineries involved. ATs bear 3 distinct domains: (i) the Sec-dependent signal sequence, that is located at the N-terminus, directs the secretion of the precursor across the IM; (ii) the central passenger domain that carries the functional site; and (iii) the carboxy terminal transporter domain, that inserts as a β-barrel into the OM and facilitates transport across the OM (10–13).

ATs function as adhesins (e.g. Ag43 of *Escherichia coli*) (14), toxins (e.g. VacA of *Helicobacter pylori*) (15), esterases (e.g. EstA of *Pseudomonas aeruginosa*) (16) and proteases (e.g. the Arginine-specific Aminopeptidase of *P. aeruginosa*, AaaA (17), and the serine protease of Enteropathogenic *E. coli* (EPEC), EspC (18)). The passenger domain of some ATs, e.g. EspC, is autoproteolytically cleaved internally and released externally (19). Other ATs, e.g. AaaA remain tethered to the outer membrane of the producing bacterium by their β-barrel, with the passenger domain exposed to the external milieu (17).

During their export to the outer membrane, several ATs have been detected at the bacterial pole (20). This localization is dependent on factors including (i) functional lipopolysaccharide (LPS) (20), (ii) the *E. coli* cytoplasmic chaperone DnaK (21), and (iii) a conserved cell division component FtsQ (22). The bacterial actin homologue (MreB) is the main component of the bacterial cytoskeleton (23, 24) which influences chromosome segregation and cell polarity in addition to maintaining cell shape (25, 26). The bacterial cytoskeleton is helical, with MreB forming cytoplasmic filaments which dynamically rotate around the cell circumference along the long axis in a manner reliant on the cell wall synthesis machinery (27). It has been suggested that the MreB cytoskeleton could act as a scaffold to provide paths for membrane protein localization and diffusion (28). In support of this, defects in MreB synthesis caused mislocalization of the type IV pilus retraction ATPase, PilT, away from bacterial poles of *P. aeruginosa* mutant strains (28), and disturbance in the localization of the AT IcsA and several polarized proteins in *Caulobacter crescentus* (29). In addition, MreB filaments have been implicated in the polar localization of aspartate chemoreceptor, Tar, leading us to speculate that polar localization of ATs could utilize MreB.

Despite lacking a typical Sec-targeting signal peptide, localization of MreB is dependent on the Sec system (30) revealing a previously unrecognized role of Sec in organizing bacterial cells on both sides of the IM. Moreover, Sec translocon proteins are distributed along the cytoskeleton in a spiral-like structure (31–34), and the secretion of Colibactin-maturating enzyme (CIbp) across the IM is MreB and Sec/SRP/YidC dependent (35). To reflect this body of literature, we extended our hypothesis to propose that MreB helical filaments in combination with the Sec machinery may act as a pathway to deliver ATs toward bacterial poles.

To investigate the localization patterns of autotransporters as they are secreted from a bacterial cell we used two separate methods of fluorescent labelling to follow ATs with proteolytic passenger domains (EspC and AaaA) as they move out of the bacterial cell. We show for the first time that AT localization is dependent on a dynamic interaction with a cellular structure reliant upon the actin homologue MreB, and the Sec translocon, and occurs in a helical distribution along the bacterial cytoskeleton.

## Results

### ATs with protease activity localize to a spiral structure resembling the bacterial cytoskeleton

To track localization during secretion, two fluorescent tags were introduced into two protease ATs. A cell-tethered AT (AaaA) and an extracellularly released AT (EspC) were selected to identify common secretion pathways. Fusion of the ATs to the monomeric variant of DsRed fluorescent protein, mCherry, provided fluorescence directly (36, 37). The other tag (i) involved fusion to a smaller polypeptide, the tetracysteine (TC) motif (two cysteine pairs flanking Proline and Glycine residues), and (ii) provided enhanced experimental control in the timing of labelling since it requires exogenous addition of the biarsenical labelling reagent FlAsH-EDT2 to become fluorescent (38). The mCherry was fused adjacent to Lys residue 340 of the EspC passenger domain (Fig. S1*A*). We predicted this site would not disrupt the EspC catalytic serine at position 256 (39). The TC-motif was genetically engineered into the EspC passenger domain between Gln530 and Phe531. The *in situ* modelled structure of EspC predicts this site is externally exposed, and distant from the active sites, and would thus be easily accessible by the FlAsH substrate without affecting the function of the target protein (Fig. S1*B*).

Fluorescence microscopy revealed that EspC-mCherry was distributed in patches. Some was at the cell poles and some distributed along the length of the cell in a defined helical arrangement (Fig. 1*A* left panel and 1*B*). Stacking individual confocal sections revealed the extended helical patterns along the length of the cell in 48% of the bacteria examined (Fig. 1C and Movie S*1*). Control bacteria producing native mCherry exhibited diffuse cytoplasmic localization throughout the cell as expected (right panel Fig. 1*A*). EspC labelled with FlAsH (EspC-TC) also acquired a spiral patchy localization (Fig. 2*A* and Fig. S2). No spiral fluorescence was observed with the unlabelled control EspC. Super resolution microscopy (SR-SIM) revealed that the 3D helix was formed from patches of protein encircling the cell wall of the bacterium (Fig. 2*B*). 3D rendering of structured illumination images allowed a panoramic inspection inside the bacterium during EspC secretion (Fig. 2*C* and Movie S2). The striking spiral distribution of EspC molecules was evident at the boundary of the membranes or within the periplasm. Patches of fluorescence were clearly apparent around the cell membranes, stretching along the bacterial cytoskeleton from pole to pole (Fig. 2*C* and Movie S2). Most notably the lack of the FlAsH labelled EspC within the bacterial cytoplasm highlighted the distinct possibility that the bacterial cytoskeleton could be driving some kind of localization during AT secretion.

**Fig. 1.**
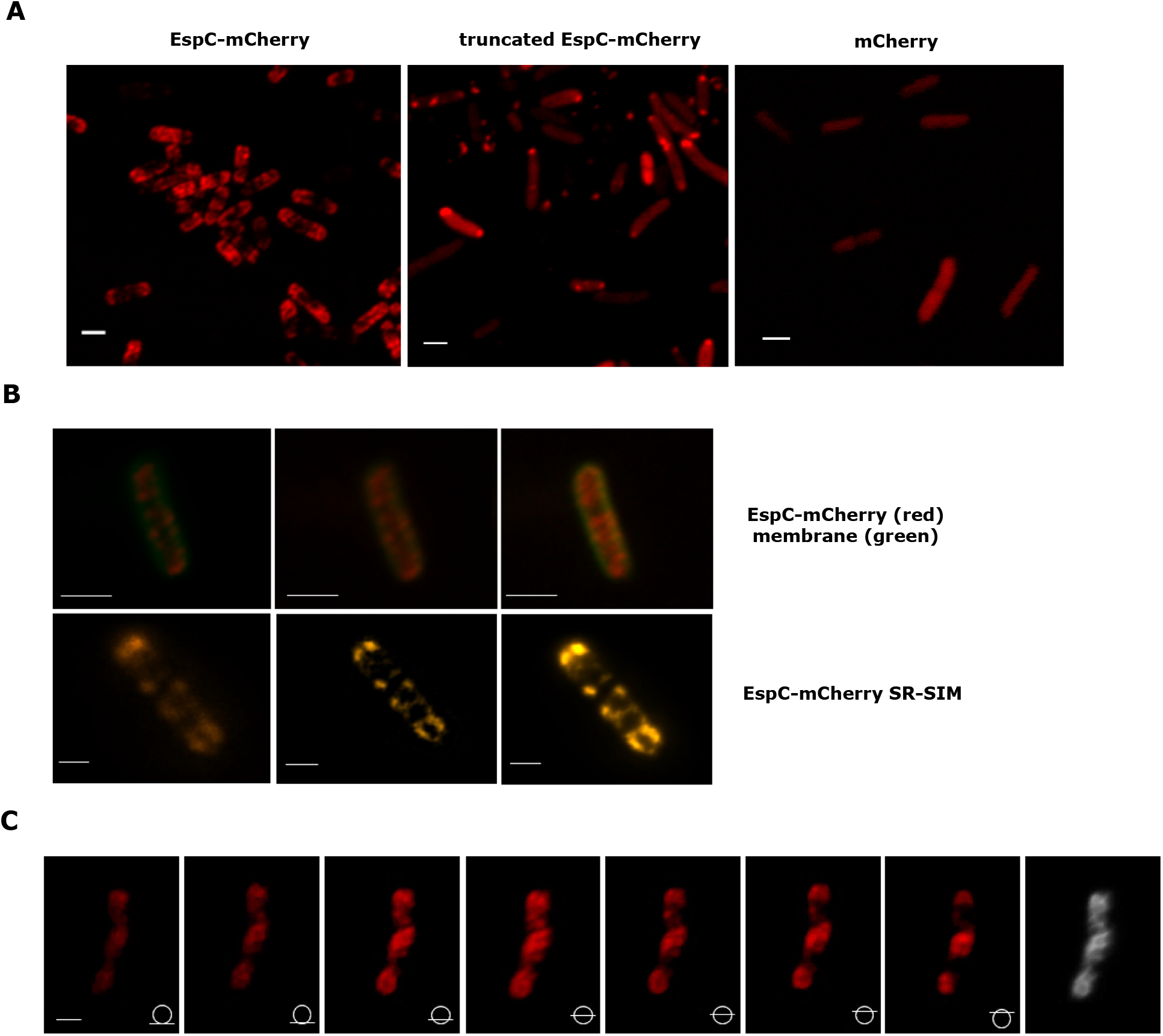
EspC locates to a spiral structure that resembles the bacterial cytoskeleton when it has its β-barrel domain attached. Bacteria were grown in LB containing 0.02% arabinose to induce mCherry containing proteins, which were imaged by fluorescence microscopy. (*A*) The left panel shows confocal imaging of *E. coli* MG1655(p33espCmcherry) producing the full length EspC-mCherry localising in a patchy helix. The central panel shows *E. coli* MG1655(p33tespCmcherry) producing truncated EspC-mCherry that does not adopt a helix localisation. The right panel shows *E. coli* MG1655(pm33Cherry) producing diffusely localized mCherry in the absence of EspC. (*B*) The position of the membrane relative to mCherry is shown in the top row by staining *E. coli* DH5α(p33espCmcherry) membranes with green FM 1-43 and imaging by confocal microscopy. In the bottom row, SR-SIM imaging of *E. coli* MG1655(p33espCmcherry) is shown to demonstrate the enhanced resolution of the EspC-mCherry helix in red. (*C*) Individual confocal images of a Z stack through a MG1655(p33espCmcherry) reveals the extended spiral fluorescent pattern adopted by full length EspC-mCherry. The diagram presented in each image (bottom right), corresponds to the position of the Z stack. Movie S1 is an animation of these images. An image with fully extended spiral structure was further processed by using ImageJ software showing the helix in grey (in the far right panel). Scale bar represents 2 μm in (*A*, top row of *B*) and 1 μm in (bottom row of *B*, *C*).

**Fig. 2.**
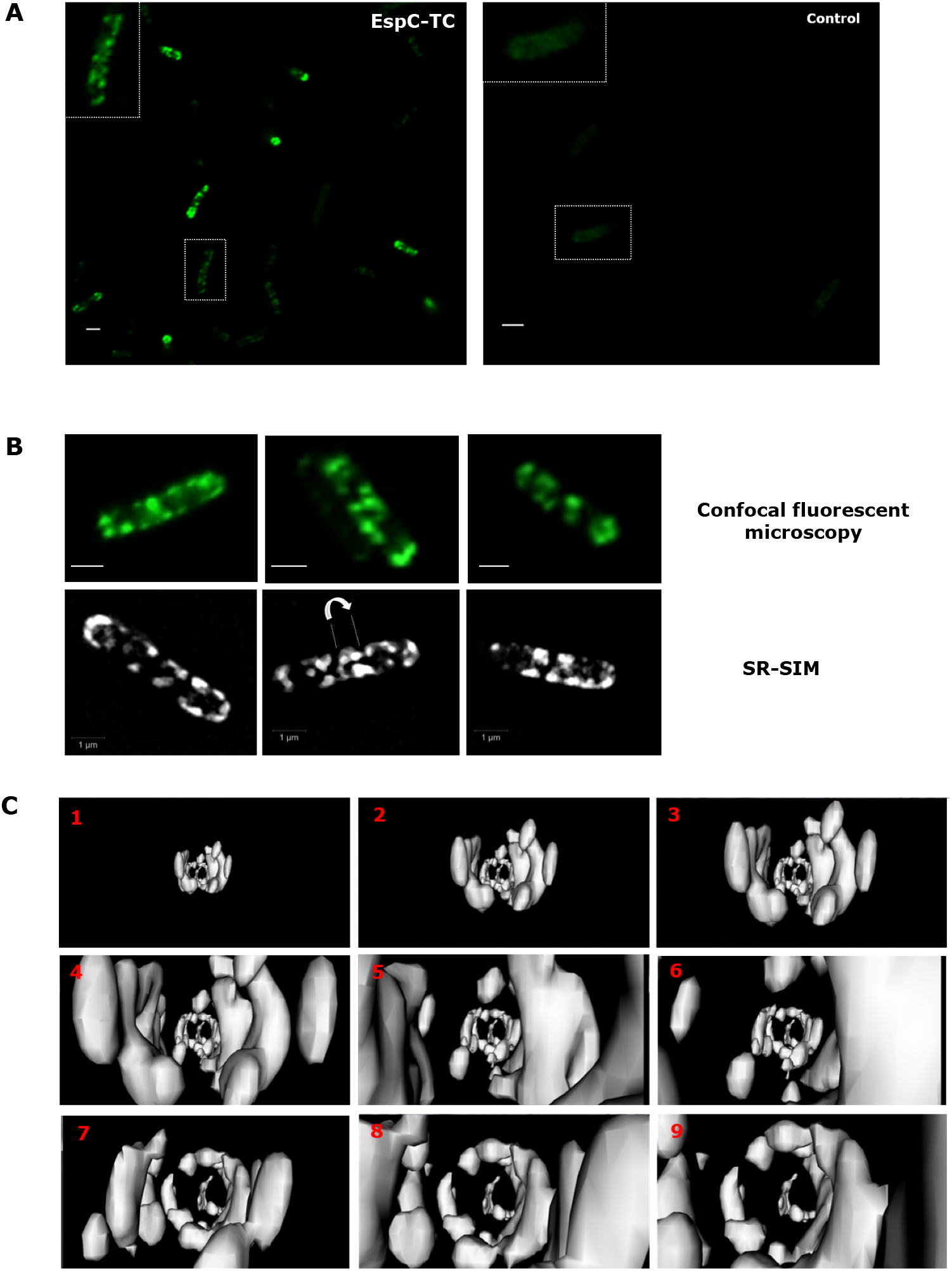
The spiral arrangement of FlAsH labelled EspC-TC aligns exclusively with the cell envelope. (*A*) MG1655(p33espC-TC) and MG1655(p33espC) were grown in LB medium, induced with 0.02% arabinose, labelled with FlAsH and imaged by fluorescence microscopy to reveal a helix for EspC-TC (left panel) or no visible structure for the control EspC (right panel). (*B*) Individual MG1655 cells producing EspC-TC were imaged by confocal microscopy (top row) and at higher resolution by SR-SIM (bottom row). The arrow in the middle picture in the bottom row indicates the diagonal pattern of EspC around the bacterial cell circumference. (*C*) 3D rendering of SR-SIM images reveal the patchy helical localization of FlAsH labelled EspC-TC exclusively within the cell envelope (longitudinal view). A collection of each frames 1-9 in sequence was depicted as a 3D animation movie in Movie *S2*. Scale bar represents 1 μm.

To test whether the observed localization represents a common underlying secretion pathway for ATs, localization of the surface exposed AT AaaA was compared with that of EspC. The TC sequence was inserted after the Leucine residue at codon 291 of AaaA to optimise fluorescence whilst minimising disruption of AT activity creating AaaA-TC. Confocal fluorescence microscopy revealed that FlAsH-labelled AaaA-TC was localized in patches orientating circumferentially around and along the long axis of the cell (Fig.3*A*). Z-stack confocal images further supported the AaaA localization as a helical array (Fig.3*B*). Control bacteria producing the AaaA that was not labelled by the FIAsH substrate showed very weak diffuse fluorescence signals (Fig.3*A*). Higher resolution SR-SIM confirmed that during secretion AaaA localized to patchy helices around the bacterial circumference very similar to those of EspC (Fig.3*C* and Movie S3).

**Fig. 3.**
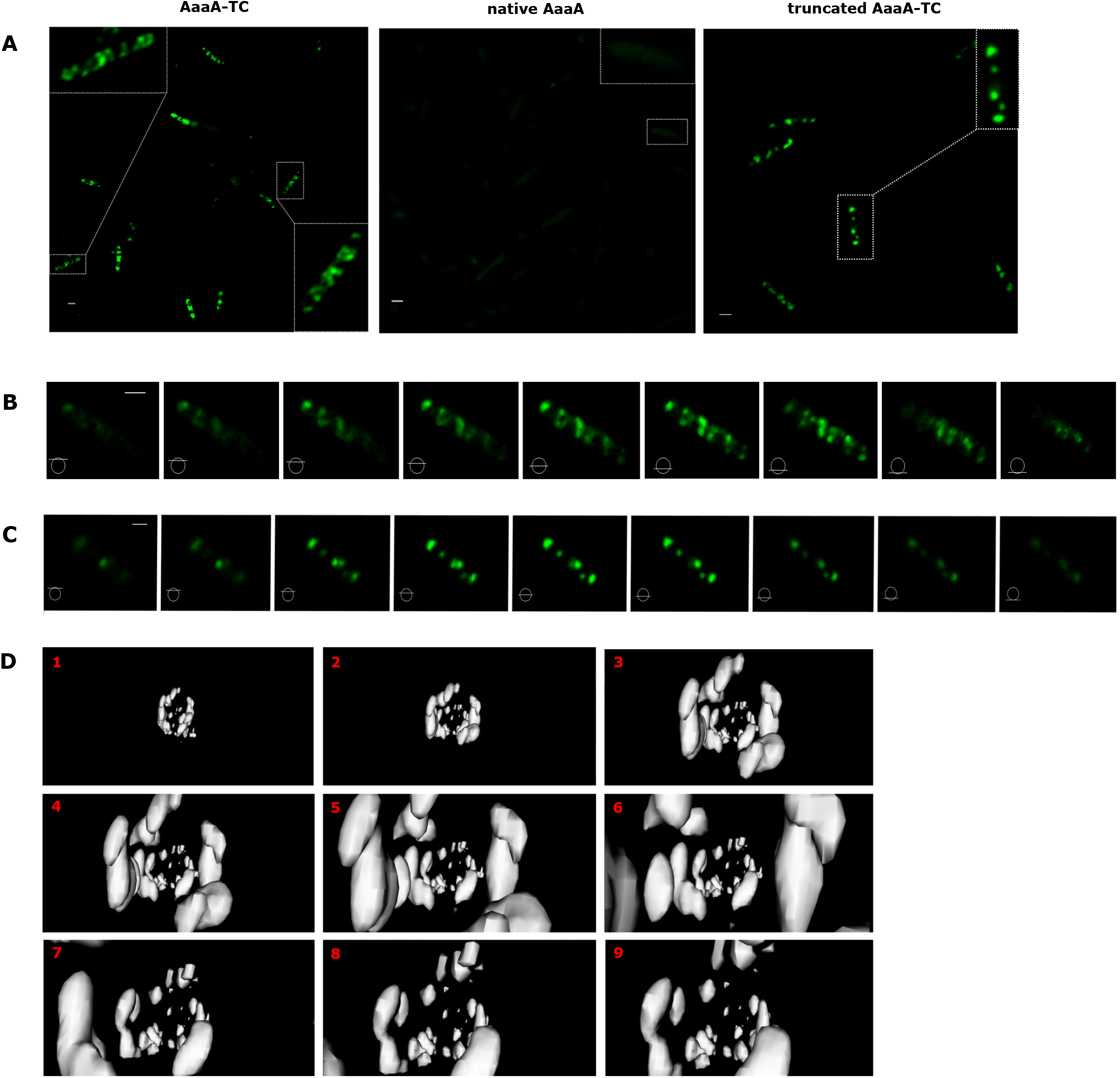
FlAsH-labelled AaaA-TC with an intact β-barrel domain adopts a helical patchy configuration. (*A*) MG1655 containing p33aaaA-TC (producing AaaA containing the TC motif), p33aaaAC233S (producing control AaaA lacking the TC-motif and native consecutive cysteine residues), or p33taaaA-TC (producing AaaA with the TC motif but lacking the C-terminal β-barrel domain) were induced by 0.02% arabinose for 2.5 h and labelled with FlAsH. Fluorescence microscopy shown here reveals cells producing FlAsH labelled AaaA-TC display a patchy helical pattern whereas those producing truncated AaaA-TC localize as a dense random distribution of fluorescent spots with no structured helical arrangement. Negative control, AaaAC233S was engineered to remove the two consecutive native cysteine residues (located at positions 232 and 233) that were able to interact with the FlAsH substrate in the absence of the TC tag, by exchanging cysteine 233 with serine by site directed mutagenesis. It was active and produced at comparable levels to AaaA. Confocal Z-stack sections of the bacteria producing TC-tagged truncated AaaA (*C*) lack the spirals observed with AaaA-TC (*B*). The diagrams of bisected circles (bottom left of each image) show where the focal plane was in the cell for each image in panels B and C. (*D*) 3D rendering of structured illumination images reveal the spiral localization of FlAsH labelled AaaA-TC within the cell envelope. A 3D animation movie was created (Movie S3), and individual frames from 1-9 are shown here. Scale bar represents 1 μm in (*A*) 2 μm in (*B*, *C*).

### Spiral formation was dependent upon the presence of the AT β-barrel

A truncated version of EspC-mCherry lacking the C-terminal β-barrel domain (produced from plasmid p33tespCmcherry) exhibited a diffuse localization with polar accumulation that lacked a helical structure in 100% of the bacteria examined (Central panel, Fig. 1*A*). Likewise, truncated AaaA-TC lacking the C-terminal β-barrel domain (AaaA-TCΔ) did not localize to a helix (Fig. 3*A* right panel). AaaA-TCΔ was engineered by introduction of an amber stop codon (TAG) at amino acid position 343 in the sequence of AaaA-TC (which is equivalent to position 338 in the native AaaA sequence). Unlike truncated EspC-mCherry, AaaA-TCΔ localized to dense foci (blobs) distributed along the long axis of the cells (Fig. 3*A* right panel). Optical sections of a bacterium producing FlAsH labelled AaaA-TCΔ confirmed the absence of the spiral localization array (Fig. 3*C*, Movie S4 and Fig. S3).

### Perturbing the actin homologue MreB alters the helical organization of ATs

To test the possibility that the bacterial cytoskeleton actin homologue MreB influences the spiral arrangement of EspC and AaaA during secretion, the localization of MreB was first tracked. To visualize the localization pattern of MreB, the TC motif was inserted by engineering *mreB* to encode two cysteine residues before and after residues Pro227-Gly228 using site-directed mutagenesis. As shown in the Fig. 4 *A* and *B*, FlAsH labelling of MG1655 producing MreB-TC showed that MreB is localized in patchy patterns. In most bacteria (62% of those examined), these patches spanned the entire length of the cell. Consistent with previous reports (40), in some bacteria (35% of those examined), short filaments diagonally followed the curve of the cells. The patchy structural pattern of MreB was particularly evident using advanced super resolution, dSTORM analysis (Fig. 4*C*).

**Fig. 4.**
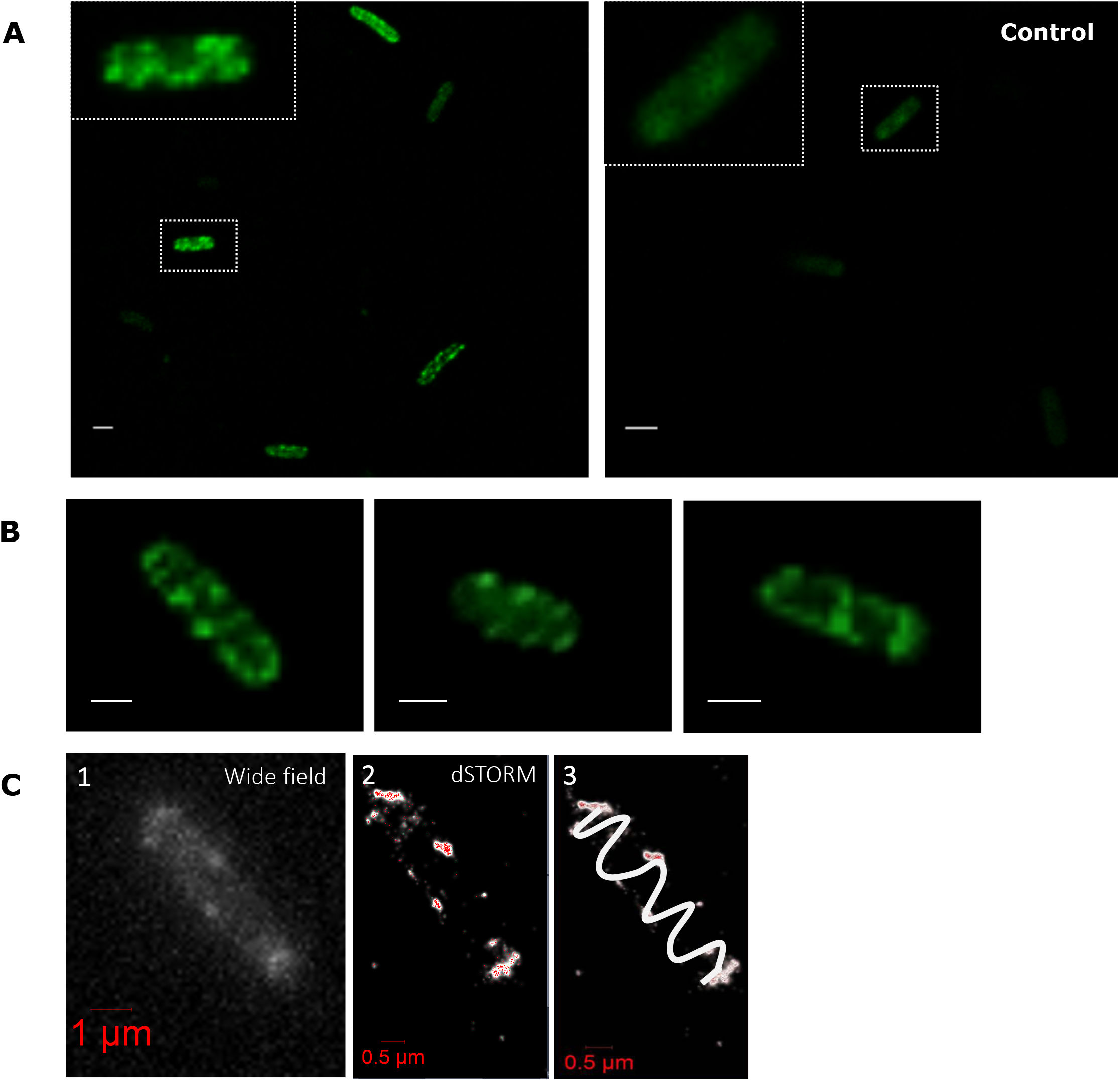
FlAsH labelled MreB adopts a patchy helical distribution. (*A*) MG1655 cells hosting either p33mreB-TC (left panel), or p33mreB (control, right panel) were induced with 0.02% arabinose for 2.5 h. Subsequently, cells were labelled with FlAsH-EDT2 and imaged by confocal microscopy to reveal helical structures for MreB-TC (left) or diffuse background staining for MreB (right). (*B*) Individual FlAsH labelled cells producing TC-tagged MreB display the patchy helical patterns of MreB with some short filaments that appear around the curve of the cell. (*C*) STORM image of FlAsH labelled MreB-TC shows the precise details of MreB localization in the bacterial cell wall (the centroids of the localized molecules in red pixels and the Gaussian rendering (visualises localization probability) of molecules in white clusters) which appears as a patchy distribution along the cell membranes (*C2*). Connecting the localized patches from pole to pole created a drawing of a spiral structure (*C3*). Scale bars indicate 1 μm in *A*, *B*.

Next, to verify the contribution of MreB to AT localization, we inhibited MreB filament assembly using the compound A22. It has been reported that the effect of A22 is similar to the effect of a MreB null mutant (41). In order to monitor EspC-mCherry localization patterns in the presence of A22, arabinose induced MG1655(p33espCmcherry) hosting the full length EspC-mCherry and MG1655(p33tespCmcherry) containing the truncated EspC-mCherry version were treated with A22^10^. As a control, a set of samples were induced and grown without treatment (A22^0^). As a consequence of using the MreB filament inhibitor A22, the morphology of the bacterial cells was changed from rod to rounded shapes as expected (Fig. 5*A*). The fluorescent localization patterns of both full length and truncated EspC-mCherry forms dramatically changed. Following treatment with A22^10^, the fluorescent localization pattern of full length EspC-mCherry was completely disrupted and appeared diffuse with no distinct spiral arrangement. In contrast, the truncated version of EspC-mCherry lacking the β-barrel was observed as strong diffuse fluorescence with irregular punctate foci (Fig. 5*B*).

**Fig. 5.**
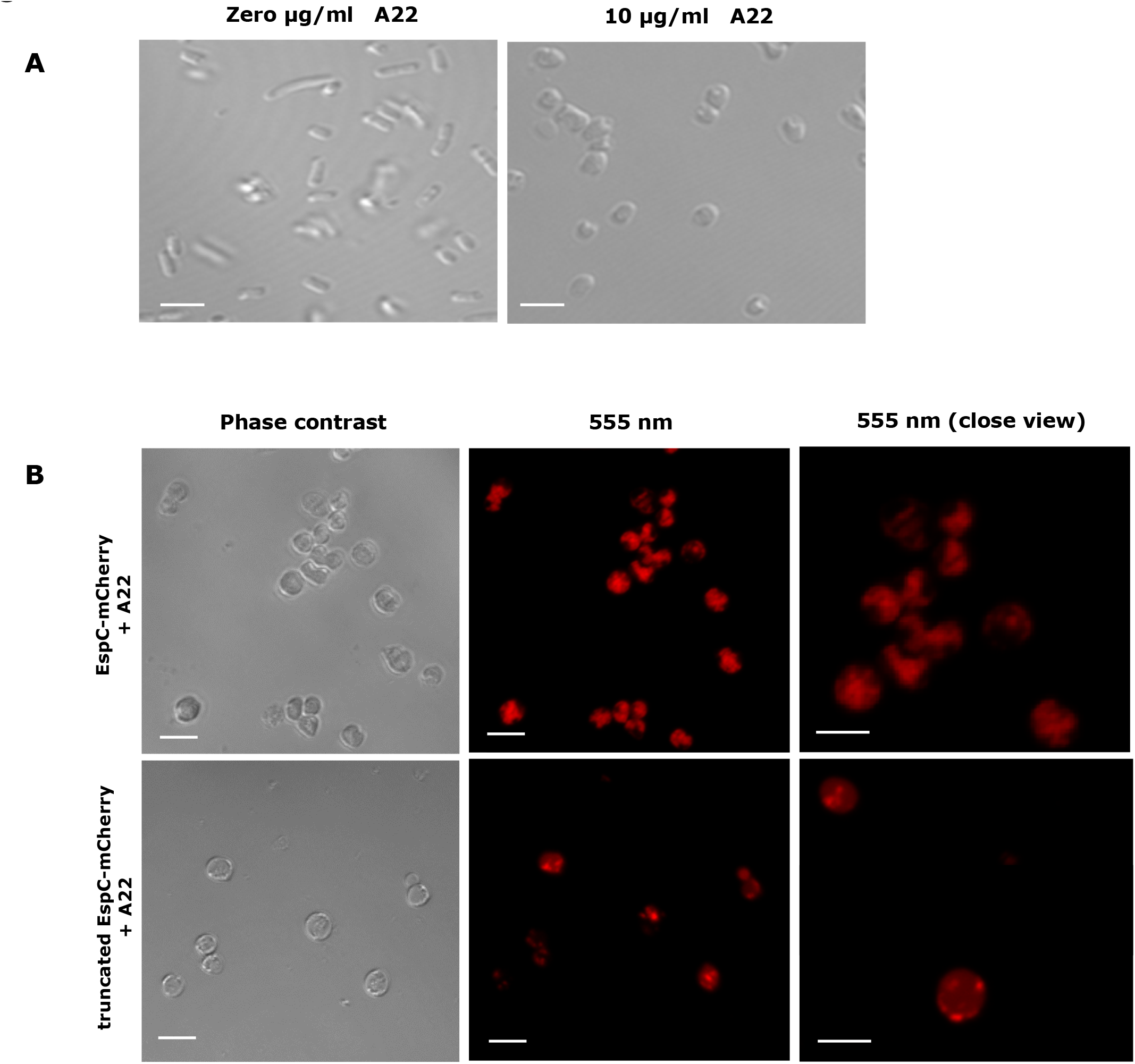
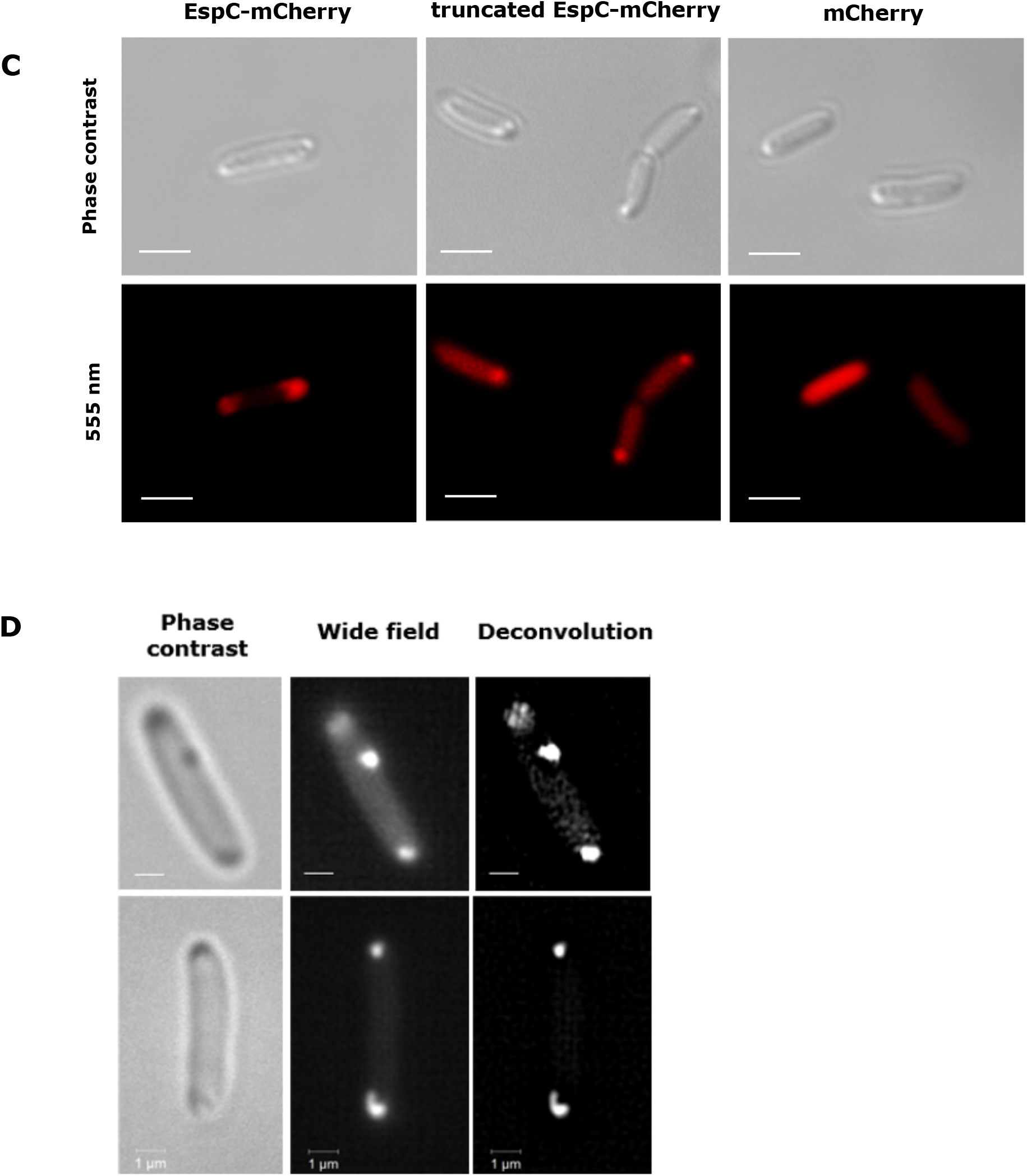
The patchy spiral localization of ATs is dependent on the bacterial cytoskeleton MreB and Sec translocon. (*A*) Inhibition of MreB filament assembly with A22 causes bacterial cell shape changes. MG1655 wild-type was treated with A22^0^ or A22^10^. Representative rounded cells treated with A22^10^ are shown following phase contrast imaging. (*B*) A22 alters the localization of EspC-mCherry with and without the β-barrel domain. 0.2% arabinose induced MG1655(p33espCmcherry) (top row) and MG1655(p33tespCmcherry) (bottom row) strains were grown and treated with A22^10^. Phase contrast and flourescence microscopy revealed full length EspC-mCherry did not form patchy spirals while truncated EspC-mCherry localized in an irregular punctate pattern in A22 treated MG1655. (*C*) The full length EspC-mCherry localization was predominantly bipolar whilst truncated EspC-mCherry was unipolar when produced by the *E. coli* SecA mutant, MM52. *E. coliΔsecA*(p33espCmcherry), *E. coli*ΔsecA(p33tespCmcherry) and *E. coliΔsecA*(p33mcherry) cultures were induced with 0.02% arabinose after 3 h of incubation at the non-permissive temperature. Phase contrast (top row) and confocal mCherry fluorescence microscopy images (bottom row) are shown. The green 555 nm laser was used to detect mCherry. (D) Loss of patchy helical localization of FlAsH labelled AaaA-TC when produced by the *E. coli* SecA mutant, MM52. Phase contrast, wide field and the more improved contrast and resolution (SR-SIM) of *E. coliΔsecA*(p33aaaA-TC) cells show the distinctive bipolar localization of arabinose induced AaaA in the absence of Sec translocon. *E. coliΔsecA*(p33aaaA-TC) was induced with 0.02% arabinose for 3 h at the non-permissive temperature. Images were taken by confocal microscopy. Scale bars represent 2 μm.

### Inactivation of SecA alters the localization of ATs

Sec has been reported to be maintained in a spiral localization (33). To investigate the intracellular localization of EspC in the absence of Sec, p33espCmcherry, p33tespCmcherry and pmLA33Cherry (control) were transformed into a SecA mutant of *E. coli* (MM52) and fluorescence microscopy was undertaken. Interestingly, both forms of EspC-mCherry exhibited localization patterns different to those observed previously. In *E. coli*ΔsecA grown at the non-permissive temperature, full length EspC-mCherry produced a bipolar and mildly diffuse localization whereas truncated EspC-mCherry was diffusely distributed or unipolar (Fig. 5*C*). Notably, the control *E. coliΔsecA*(pmLA33Cherry) exhibited the same diffuse patterns of fluorescence as observed before (Fig. 5*C*). The bipolar distribution of ATs in the absence of SecA was even more pronounced when the localization of AaaA was tracked in *E. coliΔsecA*(p33aaaA-TC) stained with FlAsH (Fig. 5*D*).

### The EspC spiral formation is unlikely to be an artefact of aggregation

Labelling EspC with the large 30 kDa mCherry fluorescent protein may interfere with its journey toward the bacterial surface and cause protein misfolding or premature folding (aggregates) resulting in fluorescence artefacts (42). To discount this, whole cell extracts induced by 0.02% arabinose from MG1655(p33espCmcherry) and the control MG1655(pBAD33) strains were subjected to floatation sucrose gradient fractionation and immunoblotting. This method was selected because it provides a good separation of membrane associated proteins from aggregated proteins (43). Importantly, the full length EspC-mCherry protein was mostly located in the fraction containing 55% sucrose which was the same fractions shown for the folded OM protein, TolC. There was also no detection of any EspC at higher sucrose concentrations (58-61%) which are associated more with aggregated proteins suggesting that the full length EspC-mCherry is associated with the OM in a folded state (Fig. 6*A*). No EspC was detected in total cell extracts from the control MG1655(pBAD33) strain (Fig. S4*A*). To further explore if different expression levels of EspC-mCherry might cause localization artefacts as a consequence of formation of aggregates, the floatation gradient analysis was repeated using whole cell extracts from MG1655(p33espCmcherry) induced with the higher (0.2%) and lower doses (0.002%) of arabinose. Similar results were obtained (Fig. S4*B*), which indicate that the EspC helix forms at different protein production levels, discounting the possibility of it being an artefact.

**Fig. 6.**
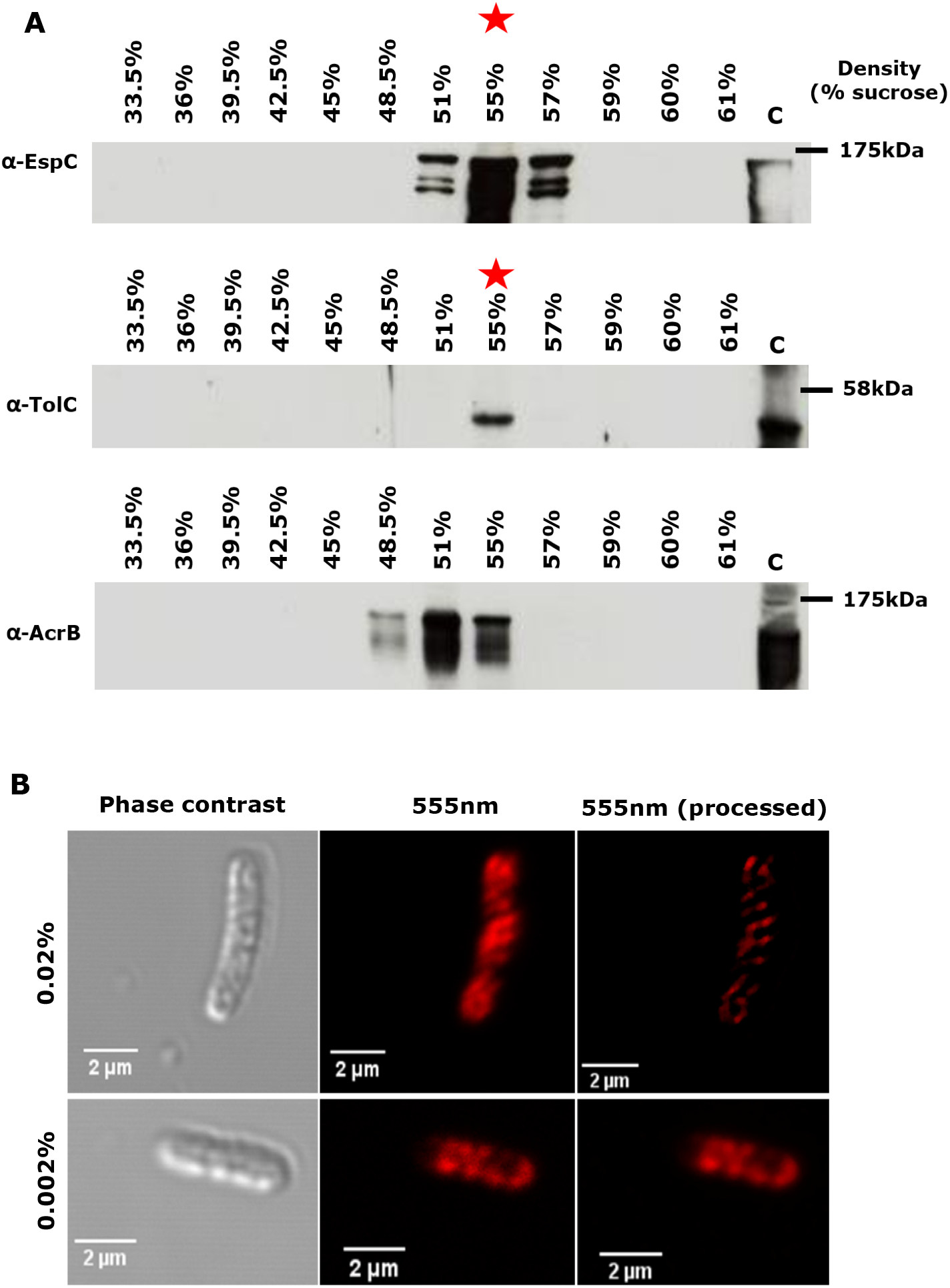
EspC-mCherry associates with bacterial outer membrane proteins and localizes to a spiral even when produced at low levels when there is a lower likelihood of aggregate formation. (*A*) MG1655(p33espCmcherry) was induced with 0.02% arabinose. A cell lysate was prepared and applied to the bottom of a 32-60% sucrose gradient. Following centrifugation, aliquots from each fraction of the gradient were separated by SDS-PAGE and immunoblotted using a-EspC (top row), the OM control a-TolC (middle row) and the IM control a-AcrB (bottom row). The sucrose concentration (% w/w) in each fraction is shown. Red asterisks indicate the location of the full length EspC-mCherry in the fraction containing 55% sucrose, which aligns to the location of TolC, an OM protein. Molecular weight markers (kDa) are shown. (*B*) Confocal images of *E. coli* cells expressing the full length EspC-mCherry (strain MG1655(p33espCmcherry)) were induced with 0.02% arabinose (top row) or with 0.002% arabinose (bottom row).

To verify that the formation of helically localized ATs was not an artefact arising from high protein production levels (37), we sought to reduce the protein production to the lowest detectable level. The microscopy data shown so far in this study was obtained using AT induction with 0.02% arabinose. Fig. 6*B* shows that culture of MG1655(p33espCmcherry) induced for 4 h with 0.002% arabinose was more than enough to observe the discrete patches and spiral patterns of EspC-mCherry inside bacteria.

### The helical localization of AT along the cytoskeleton is dynamic and transient

The circumferentially distributed disconnected patches of both EspC and AaaA hint at a dynamic spiral path being followed by the ATs during secretion. To assess the potential dynamic nature of these patches, fluorescence recovery after photobleaching (FRAP) was used in live cells producing full length EspC-mCherry fusion to determine if the localization of molecules is transient. As a control, AaaA-TC was also exploited in this experiment due to the different fluorescent labelling technique that requires addition of a substrate (FlAsH-EDT2) to produce fluorescence. MG1655(p33espCmcherry) was grown, induced with 0.02% arabinose for 4 h and prepared for confocal imaging as previously described. As shown in Fig. 7*A*, a selected single fluorescent patch (green circle) located close to the middle of the bacterium was photobleached using a focused laser beam and the recovery of fluorescence was monitored for up to 35 min. Interestingly, the EspC-mCherry patch showed a good recovery of fluorescence over time (Fig. 7*C*). The Zeiss FRAP analysis revealed that the percentage of mobile and static molecules was 67.93% and 32.07% respectively inside the green circle.

**Fig. 7.**
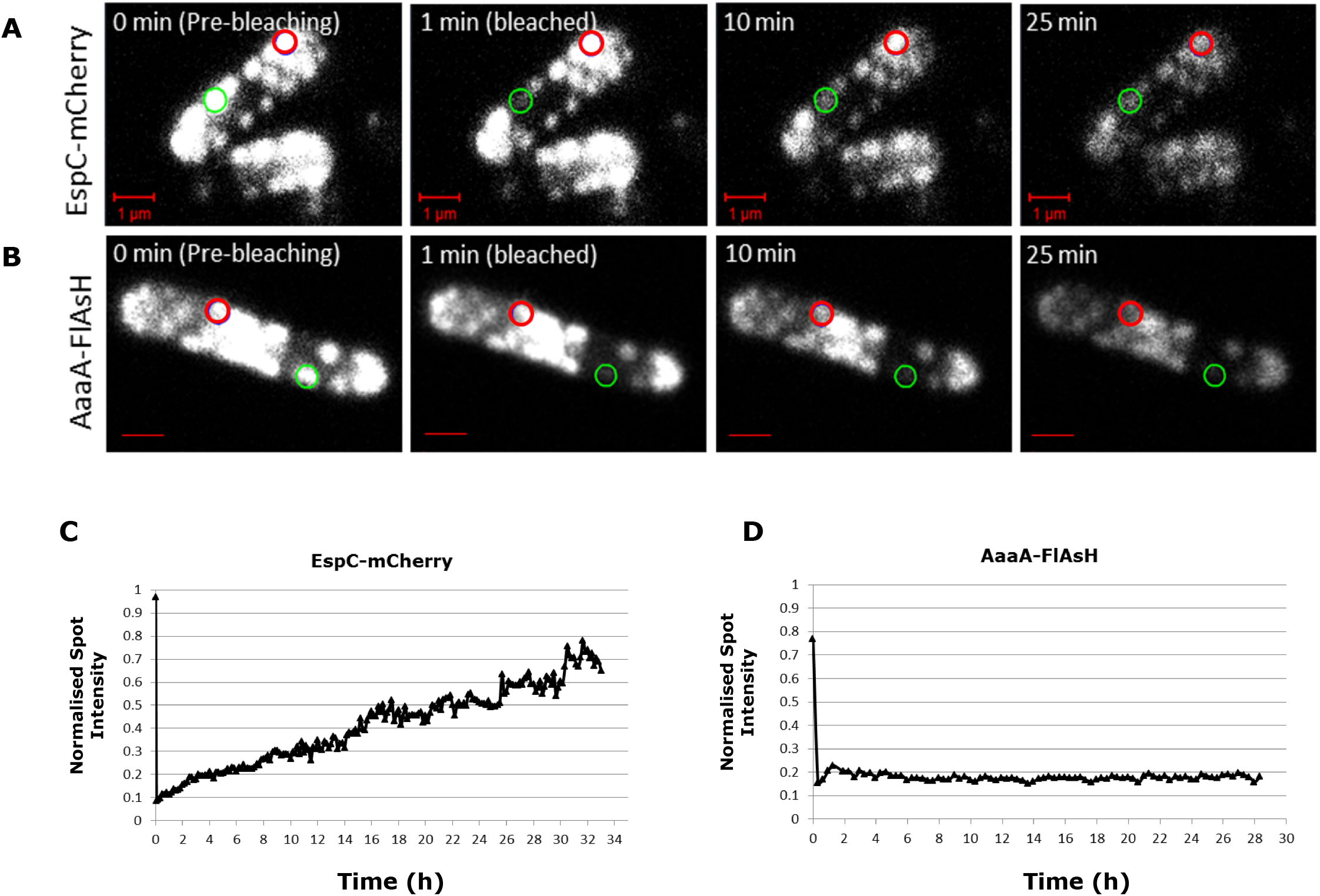
Fluorescence recovery after photobleaching reveals the dynamic nature of the molecules of EspC in patches. MG1655(p33espCmcherry) was induced with arabinose and visualised by super resolution microscopy to show the helical distribution of EspC-mCherry (*A*). MG1655(p33aaaA-TC) producing AaaA (*B*) was used in this experiment as a control, following induction with 0.02% arabinose for 2.5 h and labelling with FlAsH substrate. The representative images of EspC-mCherry (*A*) and AaaA-FlAsH (*B*) show fluorescent intensity before and after photobleaching at the region of interest marked by the green circle. Red circles indicate the control unbleached regions. The intensity of fluorescence within the red circles was used as a reference to normalise fluorescent values produced from photobleaching within the green circles accordingly. The Black triangles in graph *C* and *D* show the normalised fluorescent intensity of zones marked by the green circles in each single frame for images in A and B respectively over time.

However, as expected, in the absence of FlAsH-EDT2 substrate bleaching one discrete patch of fluorescent AaaA-TC on the cell did not show any fluorescence recovery over time (Fig. 7*B* and *D*). Here, the percentage of mobile and static molecules was 0.4% and 99.6% respectively inside the green circle. These results support the notion that the EspC-mCherry molecules are mobile, and when newly synthesized they are able to enter pathways of secretion that are visualized as fluorescent patches in this study.

## Discussion

Our data show that the cytoskeleton created by the bacterial actin homologue, MreB, could act with the Sec translocon as a scaffold to provide a pathway for membrane protein localization and diffusion. Our findings demonstrate this pathway is utilized by pivotal bacterial virulence factors (the ATs). It is a relatively recent notion that bacteria have a sophisticated spatial organization. Consequently, the underlying mechanisms integral to the formation and function of bacterial subcellular compartmentalisation remain to be elucidated in full. It is clear that the components impact upon many fundamental cellular process including ones with parallels across eukaryotes and prokaryotes such as the general secretion system and the cytoskeleton. The ability of MreB, like actin, to assemble and disassemble from filaments could provide a hitherto elusive propulsive force that enables ATs to traverse from the IM to the OM for secretion in the absence of ATP (10), and is substantiated by our observation that the interaction is dynamic and transient. MreB filaments assemble with MreC, MreD, PBP2 and RodA. This assembly is dependent on the presence of the FtsZ ring component of the cell division apparatus (23, 44), and the treadmilling motion generated by FtsZ could add a source of propulsive energy. In *E. coli*, MreB filaments are connected directly with the IM through an interaction between the MreB N-terminal amphipathic helix and the cytoplasmic part of the RodZ ring creating antiparallel double helical filaments (44, 45). There is also now evidence that links with the Sec translocon providing a mechanism to integrate organization on both sides of the IM (30), and a potential route for the ATs to dock with BAM or TAM in the OM. One potential mechanism could be targeting of the Sec translocon by the AT towards the mid-point of the cell followed by propulsion towards the poles by a mechanism driven by MreB providing time to collide with the BAM and/or TAM and subsequent translocation across the OM and final secretion close to the poles (Fig. S5).

Originally proposed as autonomous proteins using the simplest secretion machinery, it is emerging that autotransporters rely on numerous components of the cell envelope for their assembly. Understanding this mechanism at a molecular level provides the potential to understand fundamental pathways central to the physiology of bacteria with which the ATs interact during their secretion, for example SecA and MreB, and offers the potential to identify novel antimicrobials targeted towards virulence factors. Interestingly, the Gram-positive cell division protein DivIVA (46) also interacts with SecA, but it is not known if this occurs directly or indirectly *in vivo*, nor what the nature of codependence is. An important next step is to demonstrate these protein interactions directly. For the AT EspC, there is also the intriguing mystery of how it might interact with the type 3 secretion machinery (T3SS) for delivery to eukaryotic cells without being secreted through the T3SS (47, 48). Could this be mediated via a co-dependence on MreB?

It was striking how the ATs were so distinctly located in the cell envelope. The use of fluorescent tags such as fluorescent proteins and fluorescently labelled antibodies in protein localization studies has had a huge impact on discovering and understanding the molecular basis of various molecules in both eukaryotic and prokaryotic cells (37). However, the use of large fluorescent tags bears the risk of causing localization artefacts since they may interfere with the protein’s function, and must be carefully controlled (37). Swulius and Jensen, 2012 used cryotomographic microscopy to illustrate that the spiral filament formation by MreB fused with yellow fluorescent protein (YFP) at its N-terminus is an artefact, whereas when MreB was tagged within an internal loop with mCherry, it showed a patchy localization pattern similar to that of native MreB (49). Utilization of monomeric rather than dimeric YFP also avoided any unnecessary inaccuracies in the interpretation of aggregation patterns of MreB (37). In this study, fusion of two ATs to mCherry or utilization of the much smaller TC motif and FlAsH substrate generated similar localization patterns and provided the added benefit of revealing the dynamic nature of the interaction between the AT and the bacterial cytoskeleton.

## Materials and Methods

### Bacterial strains, plasmids and growth conditions

The bacterial strains used in this study (Table S1) were routinely grown on Lysogeny-Broth (LB) agar or liquid, shaking at 37°C with chloramphenicol (25 μg/ml), ampicillin (100 μg/ml) or streptomycin (100 μg/ml) where appropriate and kept frozen at -80°C in 25% (v/v) glycerol for future use. Bacterial growth was monitored by estimating optical densities at 600 nm. *E. coli* ΔSecA strains were grown at 30°C up to OD600 of 0.5 and then switched to the non-permissive temperature of 42°C. Recombinant proteins were produced for 4 h with 0.2% or 0.02% (w/v) L-arabinose for 4 h shaking at 37°C, unless otherwise stated. All plasmids used in this study are listed in supplementary Table S2.

### DNA manipulation and cloning

Oligonucleotide primers (Table S3) were designed using American SnapGene^®^ software and were manufactured by Sigma-Genosys. Plasmid DNA was purified using a plasmid purification kit (Qiagen) or GenElute™ Plasmid Miniprep Kit (Sigma). Restriction enzyme digestion, ligation and agarose gel electrophoresis were performed using standard methods (50). Routinely, PCR reactions were performed using the Q5^®^ High-Fidelity 2x Master Mix kit (NEB) following the manufacturer’s instructions. DNA products were separated by agarose gel electrophoresis in 1x tris-acetate-ethylenediaminetetraacetic acid (TAE) buffer. DNA was purified from agarose gels using Gel extraction kit (Qiagen), or directly using a PCR purification kit (Qiagen). Plasmids were introduced into freshly prepared CaCl2 competent cells by heat shock at 42°C for 4 min. When necessary, DNA sequencing was conducted by Source BioScience, Nottingham, UK. Alignment of DNA or protein sequences to identify homology was achieved using the Basic Local Alignment Search Tool ‘BLAST’ provided by NCBI (http://blast.ncbi.nlm.nih.gov/Blast.cgi). The Sequences of DNA and protein were obtained from the NCBI (http://www.ncbi.nlm.nih.gov/) using gene and protein databases respectively. The plasmids used or generated in this study are listed in Table S2, the primers used in Table S3. Levels and activities of the recombinant ATs are shown in Fig. S6-S7.

### Site directed mutagenesis

Site-directed mutagenesis was carried out by the QuikChange II Site-Directed Mutagenesis Kit, according to the manufacturer’s instruction manual (from Agilent Technology). Both forward and reverse primers (Table S3) were designed by SnapGene software to insert or substitute particular nucleotides on desired fragments to generate mutations. Mutation was confirmed by sequencing using primers listed in Table S3.

### Inhibition of MreB using S-(3.4-Dichlorobenzyl)-isothiourea (A22)

To perturb the bacterial actin homologue cytoskeleton MreB, A22 (Calbiochem) was used (41). A22 was prepared at a concentration of 5 mM (stock concentration) and used either at zero μg/ml (A22^0^) as untreated controls or 10 μg/ml (A22^10^) for MreB inhibition. Bacterial strains were grown o/n in LB broth plus appropriate antibiotics, diluted into fresh LB plus appropriate antibiotics, induced with 0.2% arabinose, and grown with shaking to an OD600nm of 0.05. Two equal aliquots of 25 ml were removed. One aliquot was treated with A22^0^ (control) whilst the other aliquot was treated with A22^10^. Both sets of cells were grown at 37°C for a further 4 h with shaking, harvested by centrifugation at 6000 rpm for 5 min, washed once with ice cold PBS, and then resuspended in PBS to 0.5 OD600 units/ml. Finally, 8 μl of washed cells were applied separately to microscopic slides for microscopy.

### Sodium dodecyl sulphate-polyacrylamide gel electrophoresis (SDS-PAGE), Immunoblotting, cellular fractionation and sucrose gradient analysis

SDS-PAGE and immunoblotting were performed as described previously (50) using 9% SDS-PAGE gels. To enable detection of EspC, AaaA, mCherry, TolC and AcrB proteins, the rabbit α-EspC was used at concentration 1: 1000, rabbit α-AaaA was used at 1:1000, rabbit α-TolC used at 1:2000 and rabbit α-AcrB used at 1:2000 respectively as primary antibodies, whereas the goat anti-rabbit IgG conjugated to horseradish peroxidase (HRP) (Sigma) was used at a dilution of 1:10,000 as secondary antibody. All antibodies were incubated for 1 h at room temperature with shaking and washed with PBST 0.5%. Sucrose gradient fractionation was undertaken as described previously (43).

### FlAsH labelling of tetracysteine-tagged recombinant proteins

To enable detection of recombinant proteins containing the tetracysteine (TC) motif, the green membrane-permeable biarsenical dye, FlAsH-EDT2 (Invitrogen), was used in this study. Cultures of the strains to be tested were prepared and induced by 0.02% arabinose unless otherwise stated. Cells producing TC-tagged protein were harvested by centrifugation at 5500 rpm for 2 min and washed once with ice cold 1x HEPES buffered saline BioUltra (Sigma), and resuspended in 1 ml ice cold 1x HEPES to OD600 of 1.0. The cells were centrifuged and gently resuspended in 500 μl 1x HEPES buffer containing 3-5 μM FlAsH-EDT_2_ substrate followed by incubation for 35-45 min in darkness at RT. After incubation, the cells were harvested by centrifugation as before and the supernatant discarded. Subsequently, cells were washed twice with 1 ml of 1x British Anti-Lewisite (BAL) wash buffer at a final concertation of 0.25 mM and then further washed once again with the 1x HEPES buffered saline, ice cold and resuspended in 1x HEPES to approximately OD600 of 0.5. Eight μl of washed cells were mounted on microscope slide, covered with coverslip, sealed with transparent nail polish and examined by fluorescence microscopy.

### Confocal fluorescence microscopy

To generate more detailed images, a Zeiss LSM 700 confocal laser scanning microscope fitted with a HXP 120C lamp and a Zeiss alpha-Plan-Apochromat 63x/1.46 Oil objective lens was used. Images were captured by using ZEN software and analysed using image J software. The lipophilic fluorescent membrane stain, FM 1-43 (Invitrogen) was used according to the manufacturer’s instructions. Briefly, cell pellets were resuspended in 1 ml ice cold PBS containing 5 μg/ml FM-1 43 (final concentration) and incubated on ice for a maximum 1 min in darkness and then washed twice with ice cold PBS and resuspended in 1 ml PBS before mounting on microscope sides.

### Fluorescence recovery after photobleaching (FRAP)

The microscope Zeiss Elyra PS.1, with an alpha-Plan-Apochromat 100x/1.46 Oil DIC M27 Elyra objective lens in confocal mode was used for FRAP experiment. Images were acquired using either a 561 nm wavelength laser at 2.0% or a 488 nm wavelength laser at 0.3% for mCherry and FlAsH-tag fluorescences, respectively. Zeiss Zen Black 2012 software was used to extract the intensity values from the bleached and control regions over time. Drift correction was applied using Zen Blue 2012 software. Values were normalized by a direct comparison of bleached over control zone values and then plotted.

### Super resolution microscopy

For super-resolution imaging, a Zeiss Elyra PS.1 microscope was used in SIM (structured illumination microscopy) or dSTORM (direct stochastic optical reconstruction microscopy) modes. In SIM mode, either Plan-Apochromat 63x/1.4 Oil DIC M27 or C-Apochromat 63x/1.2W Korr M27 objective lenses were used. For FlAsH labelled bacteria, a filter set LBF-488/561, laser power 488 nm at 4.0% and cmos camera exposure time 25.0 ms was used in SIM mode whilst a BP 570-620+LP 750 filter, 561 nm laser at 2.1% and camera exposure 100.0 ms was used for mCherry expressing bacteria. For the dSTORM mode, an alpha Plan-Apochromat 100x/1.46 Oil DIC M27 Elyra objective, BP 495-550+LP 750 filter 488 nm laser at 100% and 405 nm laser at 1%, 100 ms exposure, EM-CCD gain 200 was used for both fluorescent mechanisms.

## Acknowledgements

We would like to thank Libyan Embassy (Cultural Attaché - London) and the School of Life Sciences postgraduate support team for financially supporting MA. This work was supported by the Biotechnology and Biological Sciences Research Council UK [grant number BB/L013827/1] (United Kingdom) which provided the means to establish the super resolution Microscopy infrastructure with which to deliver the analysis. We also thank colleagues at the University of Nottingham Centre for Biomolecular Sciences for supporting this study, in particular Louise Arnold and Stephanie Pommier for making plasmids p33espC and p33espCmcherry respectively, and to Mathulah Sriskandarajah for performing some of the AaaA activity assays.

## Author contributions

Mahmoud Ashawesh generated all the data in this study working closely with Robert Markus to produce and analyse high quality images on the super resolution microscope. Kim Hardie conceived, directed and wrote the manuscript with input from Mahmoud Ashawesh and Christopher Penfold.

